# The evolution of cavefish odor perception shifts fish response from avoidance to approach when exposed to alarm and death odors

**DOI:** 10.1101/2025.10.01.679817

**Authors:** Zachary Harkinish-Murray, Tiffany Chan, Jerry N Gbarlea, Agniva Sinha, Anthony Bedard, Hala AlGarni, Desiree T Sukhram, Javier F Juarez, Roberto Rodriquez-Morales, Robert A Kozol

## Abstract

An organism’s survival is dependent on activating the correct behavioral circuit in response to a stimulus. While stimulus perception is largely conserved, we still do not know the evolutionary mechanisms that drive changes in stimulus perception. Therefore, we wondered how odor perception has evolved in cavefish that encounter vastly different odors compared to their surface ancestors. While surface populations show attraction to social odors and avoidance of alarm and death odors, cave populations are sexually dimorphic in their responses to social odors and exhibit attraction to alarm and death odors. Finally, we find that odor perception variation is genetically encoded and plastic, while tissue clearing and brain mapping suggest odor perception is governed by thalamic and hypothalamic circuits. Taken together, we describe a unique model for investigating the environmental, genetic, and neurological mechanisms underlying olfactory evolution in vertebrates.

## Introduction

Vertebrate evolution has maintained conserved sensory percepts over hundreds of millions of years due to the stability of environmental sensory signals (1, 2). Common sensory signals are received and integrated to compute the appropriate behavioral output to find food, avoid predation, find mates and overall increase the probability of survival (1, 3). While our general knowledge regarding sensory integration and behavioral output is quite comprehensive, less is known about how genetic diversity and environmental variability drive sensory perception (4–6). This includes how sensory perception of specific environmental cues are changed from positive to negative valences or vice versa.

Taxis towards, or away from, chemical signals appeared very early in the history of life, predating the Bacteria-Archaea separation, and has remained a mainstay of mobile life forms since, including in humans (7, 8). Common odors in our environment are generally characterized by our brain as being positive or negative (9, 10). For instance, death odors are released during decomposition, providing negative cues that signal danger, area avoidance and life preservation (11–13). In contrast, the smell of prey provides positive cues that trigger approach behaviors associated with foraging (14–16). While these odor percepts are generally conserved across taxa, changes in environmental pressures and life history traits can influence the evolution of odor perception (3, 13). This is increasingly relevant in the context of the anthropogenic impact on ecological niches over time, coupled to the remarkable plasticity of the olfactory system, especially in aquatic animals (3, 17). Therefore, species adapted to extreme environments with variation in food availability (18), predation (19), and social structures (20), provide unique models for understanding the biological mechanisms underlying evolution of odor perception and the adaptability of odor stimulated behavior.

An example of this is the blind Mexican cavefish, *Astyanax mexicanus*, a teleost fish species comprised of ancestral, river dwelling surface populations and over 30 cave-adapted populations of blind cavefish, of which some have independently evolved and share convergent troglobitic features (21, 22) (Fig. 1a). These include regressive traits, such as eye degeneration and loss of pigmentation, and constructive traits, including expansions of taste receptors, lateral line and olfactory epithelia (23–27). These constructive traits are thought to compensate for eye regression, with the olfactory system of cavefish exhibiting larger sensory organs and greater sensitivity to specific amino acids that may improve survival in the dark (25, 28). Despite insights on smell ability, less is known about how odor perception evolved in cavefish and whether common environmental odors evoke the same approach and avoidance behaviors as in ancestral surface fish.

**Fig. 1.**
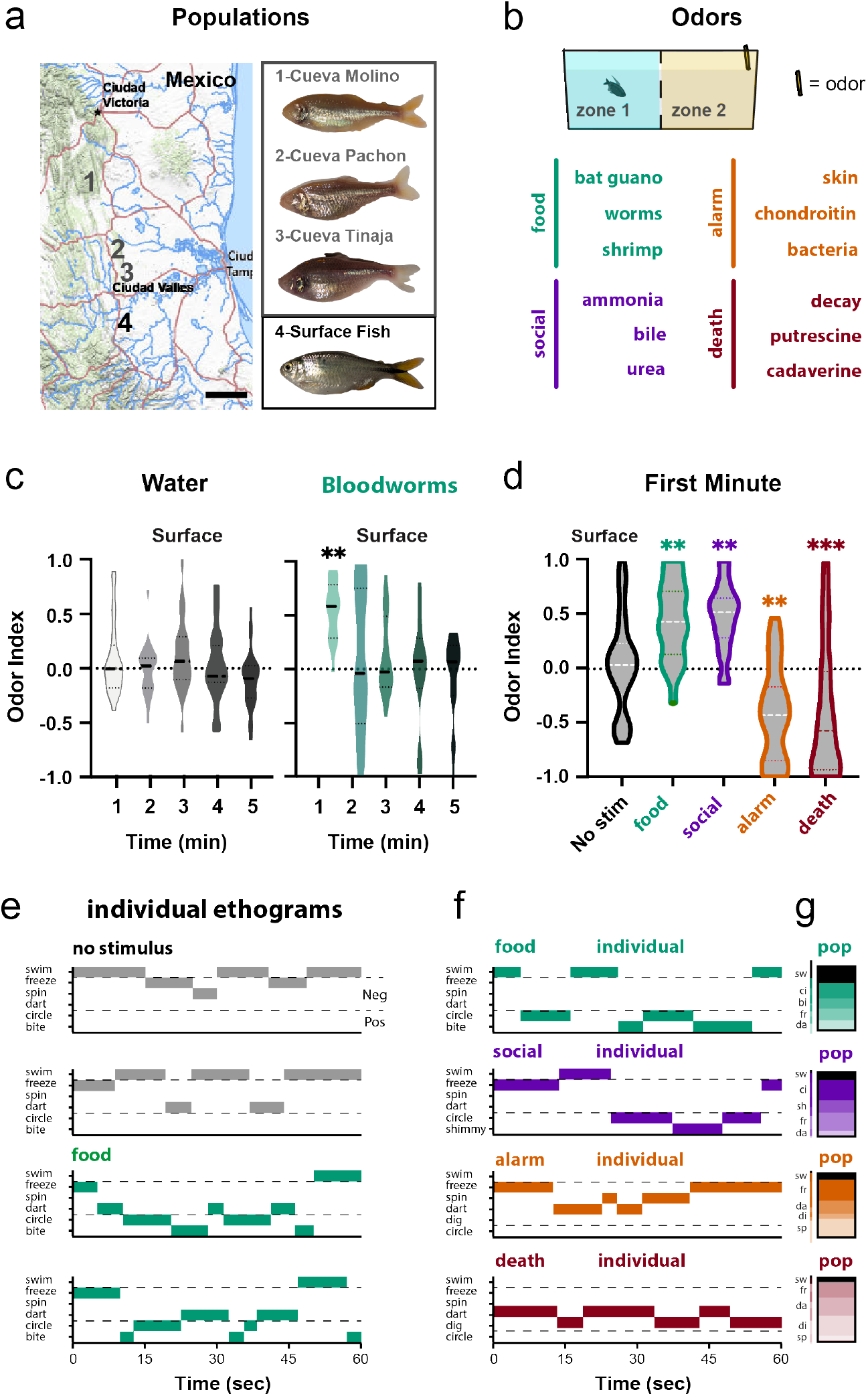
Surface fish exhibit approach to food and social odors, while avoiding alarm and death odors. a. Cavefish populations are found across the Sierra de El Abra mountain range of Northeast Mexico. b. Odors were categorized by behavioral associations in previous fish literature: food (teal), social (purple), alarm (orange) and death associated (chrimson) odors. c. Surface fish did not exhibit approach or avoidance to water control scent (n=23), but did show significant approach to the addition of water soaked in bloodworms (food scent; n=16). d. Surface fish exhibit positive approach to food and social odors (teal and purple; n=24 and n=20), while displaying negative avoidance to alarm and death odors (orange and crimson; n=27 and n=24). Representational (ef) and population (g) ethograms for food (teal; bloodworm), social (purple; ammonia), alarm (orange; skin extract) and death (chrimson; decay extract) elicited swimming behaviors. Dotted lines denote neutral (swim, sw), negative (dart, da; dig, di; freeze, fr; spin, sp) and positive (bite, bi; circle, ci; shimmy, sh) swim bouts. Indices were compared using non-parametric ANOVA’s, followed by Tukey corrected p-value multiple comparisons. p-values; *-p<0.05*, ***-p<0.01***, - p<0.001.

Here, we asked whether cavefish odor perception is variable, and if so, whether these variations impact odor stimulated behaviors across surface and cave populations of *A. mexicanus*. First, we show that surface fish exhibit similar odor perception valences (negative and positive) to zebrafish. Second, we demonstrate that light influences cavefish olfactory behavior, with darkness revealing light masked changes in odor perception. Third, we show that cavefish exhibit positive bout poses and approach behavior to odors associated with fish injury and death, suggesting a change in odor perception coding. Fourth, cavefish females and not males exhibit positive associations with social odors, suggesting that social olfaction is sexually dimorphic in cavefish. Fifth, we utilize surface to cave F_2_ hybrids to show that social, alarm and death related odor perception is heritable, which provides a functional model to investigate the genetic architecture that underlies vertebrate odor perception. Finally, we conducted tissue clearing and brain mapping to show that cavefish exhibit overlap between brain activity in the thalamus and preoptic region when exposed to food and death odors. Overall, this study reveals changes in cavefish odor perception that may underlie an improved foraging strategy in an environment with variable resources.

## Results

### Surface fish approach food and social odors, but avoid odors of injury and death

The olfactory system has evolved to discriminate between different categories of odorants, including food, social, predatory and death-associated odors (12, 29–33) To determine how odor perception has evolved in cavefish, we developed an odor discrimination assay that exposes adult fish to different odor classes dissolved in 0.5% DMSO water (Fig. 1b, Table 1). DMSO was leveraged as an anchor to maintain odor position and a biological carrier to amplify the smell (Supplementary Figure S1)(34, 35). Behavior tanks were then split into an odor zone and a no-odor zone and odor perception was then analyzed by converting the duration of zone times into an odor index, (OR=[zone 2-zone 1]/(zone 2+zone 1)), that ranged from +1.0 to -1.0 (Fig. 1b, Methods Section ‘approach and avoidance analysis’). To validate this approach, we first exposed surface fish to water shams and food-soaked water, and analyzed odor indices in 1-minute intervals (Fig. 1c&d). Using this methodology, we found that surface fish exhibit no change in behavior when exposed to a water sham and exhibit positive odor indices during the first minute of exposure to food extract (Fig. 1c, Supplemental Table S1-3). Overall, a 1-minute time bin produced positive values for the smell of worms (food) and bile (social) odors, and negative values for alarm (skin) and death-associated odors (decay) (Fig. 1d, Supplemental Table S4-5). To better contextualize the behaviors underlying these indices, we analyzed bout movements and constructed subject and population ethograms for each odor (Fig. 1e&f, Table 2). We found that surface fish exhibited swim bouts we term ‘positive’, including quick circles and biting behavior when exposed to food odors (Fig. 1e-g, Fig. 2, Supplementary Videos S1-2), and a few instances of dance-like shimmy’s when exposed to social behaviors (Fig. 1fg, Fig. 2, Supplementary Videos S3). In contrast, we found that alarm and death associated odors elicited bouts we termed ‘negative’, included freezing, darting, spinning and corner digging (Fig. 1f&g, Fig. 2, Supplementary Videos S4-7). These results clearly demonstrate that surface fish maintain ancestrally conserved approach behaviors via positive bouts (e.g. circling and biting; Fig. 1e) and avoidance behaviors via negative bouts (e.g. freezing and digging; Fig. 1e).

**Table 1.** Odorants used in olfaction experiments.

| Odorant | Odor | Ancestral Valence | Molarity (mM) | Company | Cat # |
| --- | --- | --- | --- | --- | --- |
| Water | N/A | Neutral | N/A | N/A | N/A |
| Allyl isothiocyanate (mustard) | Pain | Negative | 50%/Vol | Sigma | W203408 |
| Blood worm | Food | Positive | N/A | Bulk Reef | 208881 |
| Shrimp | Food | Positive | N/A | Bulk Reef | 212246 |
| Bat Guano | Food | Positive | N/A | Down-to-earth | 7-3-1 |
| Taurocholic acid (bile) | Social | Positive | 10 mM | Sigma | S0900000 |
| Ammonium chloride (ammonia) | Social | Positive | 40 mM | Sigma | A9434 |
| Prostaglandin F2 (hormone) | Social | Positive | 1 mM | Sigma | P5069 |
| Urea (urine) | Social | Positive | 40 mM | Sigma | U5378 |
| Skin extract (alarm substance) | Alarm | Negative | N/A | N/A | N/A |
| Pyridine-n-oxide | Alarm | Negative | 20 mM | Sigma | 131652 |
| Chondroitin sulfate | Alarm | Negative | 20 mM | Sigma | 1133570 |
| Bacteria (*Staphylococcus saprophyticus*) | Alarm | Negative | 10%/Vol | ATCC | 15305 |
| Spermine | Alarm | Negative | 20 mM | Sigma | 55513 |
| Diaminohexane | Death | Negative | 10 mM | Sigma | 613533 |
| Decay extract (death substance) | Death | Negative | N/A | N/A | N/A |
| Cadaverine | Death | Negative | 10 mM | Sigma | D22606 |
| Putrescine | Death | Negative | 10 mM | Sigma | 51799 |

**Table 2.** Behavioral bout definitions and characteristics.

| Bout Name | Definition | Tank Position | Surface Fish | Cavefish |
| --- | --- | --- | --- | --- |
| Swim (Vid S1 &S2) | Slow swim across tank length. | column | present | present |
| Freeze (Vid S3) | Immobility for longer than 5 seconds. | bottom | present | absent |
| Spin (Vid S4) | Upside down spinning on nose, back and forth. | bottom | present | absent |
| Dart (Vid S5) | Fast swim sprints back and forth. | bottom | present | absent |
| Dig (Video S6) | Driving nose into tank edge. | bottom | present | absent |
| Circle (Video S7&8) | No midline cross, turns back towards current zone | column | present | present |
| Bite (Video S9&10) | Jaw movement towards tank bottom. | bottom (bottom or column, cave) | present | present |
| Shimmy (Video S11&12) | Slow vertical movement with body shaking | column | present | present |

**Fig. 2.**
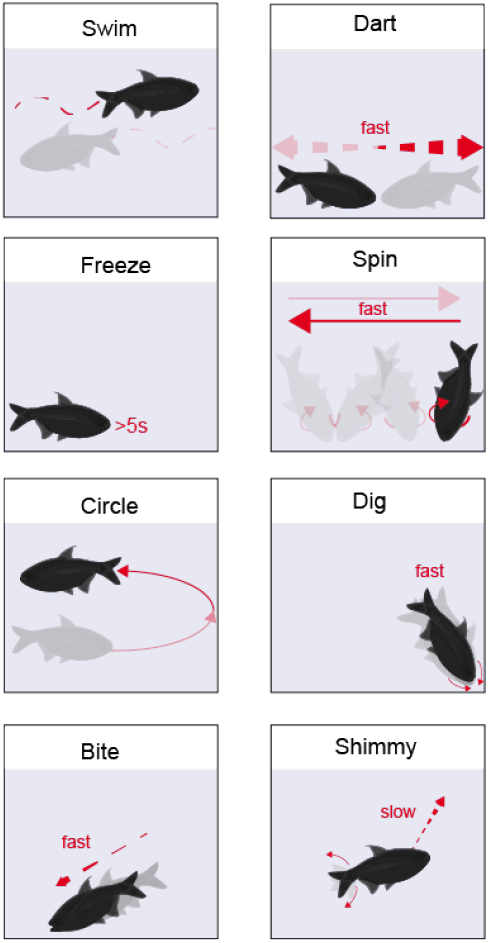
Behavioral pose pictographs for each bout type. Fish depicting the beginning of a bout are transparent and end of bout (black). Red arrows depict x,y direction and speed (fast/slow) that occurs during the bout. Positive swim bouts associated with approach are swim, circle, bite and shimmy, while negative swim bouts associated with avoidance are dart, freeze, spin and dig. All bouts were observed and defined in surface fish, while cavefish populations have lost all negative bouts.

### Cavefish exhibit positive odor indices when exposed to skin extract in the dark

Work in alarm behavior has identified several molecules released from fish skin that triggers anxiolytic responses (29, 31– 33). Previous research has shown that cave morphs of *A. mexicanus* do not perform an alarm response when exposed to skin extract from either surface or cave morphs (36). However, those experiments did not test odor responses in different lighting conditions and previous research suggests that cavefish behaviors are influenced by the presence of light (37, 38). To test whether cavefish react to alarm substances, we exposed surface fish and Pachon, Molino and Tinaja populations of cavefish to skin extract in three different light conditions: direct diffused white light, indirect white light and diffused infrared light (IR) (Fig. 3). As previously reported, cavefish showed no change in behavior when exposed to skin extract in direct diffused white light (Fig. 3b, Supplementary Table S67). In contrast, when we measured the odor index of cavefish in indirect low light and the dark, we found odor indices became positive in low light and robustly positive in complete darkness (Fig. 3e&h, Supplementary Table S8-10), suggesting that light exposure greatly influences reactivity to odor cues. At a higher level of behavioral resolution, surface fish exhibited a decrease in immobility when going from light to dark (less freezing) and increased darting time, while cavefish exhibited an increase in odor zone circling (Supplementary Videos S8), which shifted odor indices more positive from light conditions to dark (Fig. 2, Fig. 3c,f&I, Supplementary Table S11-13). Overall, these experiments demonstrate that cavefish odor perception is impacted by light and that cavefish exhibit approach behaviors when exposed to skin extract in darkness.

**Fig. 3.**
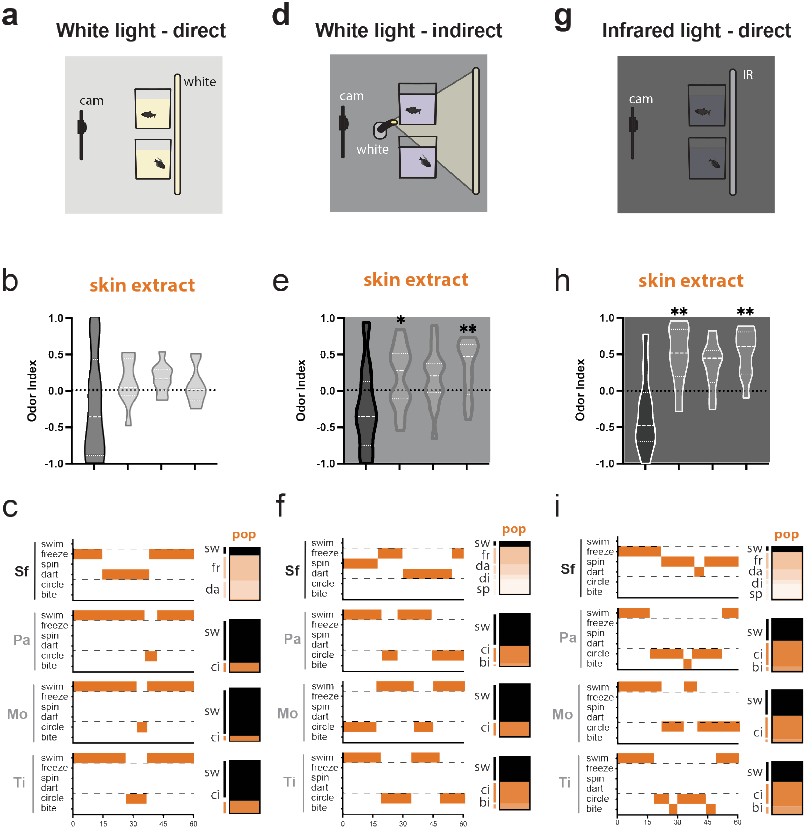
Light masks skin extract approach behaviors in cavefish. a-c. Direct white light via acrylic diffusion revealed no difference between populations when exposed to skin extract (n=9, all populations). b. Box plots reveal negative odor indices in surface fish and neutral odor indices in cavefish. c. Ethograms reveal darting (da) and freezing (fr) in surface fish, while cavefish mostly perform routine swims. d-f Indirect white light via low lux projections on a back wall revealed (e) positive odor indices in cavefish, (f) an increase in cavefish circling behavior and the emergence of head spinning (sp) and digging (di) in surface fish (sfpa, n=15; moti, n=12). g-i. Direct IR-light via acrylic diffusion resulted in (h) higher positive odor indices in cavefish and (i) revealed biting (bi) behaviors in cavefish (n=10, all populations). Indices were compared using non-parametric ANOVA’s, followed by Tukey corrected p-value multiple comparisons. p-values; *-p<0.05*, ***-p<0.01***, -p<0.001.

### Cavefish maintain ancestral odor perceptions to avoid caustic odorants and approach food odorants

To test odor perception more broadly, we performed the same odor index assay using several food, social, alarm and death-associated odors that elicit ancestrally derived reactions in other fish taxa (12, 29–33). First, we tested water shams as a neutral odor and found that neither surface or cavefish populations reacted to a small jet of water (Fig. 4a&b, Supplemental Figure S2, Supplemental Table S14). Next, we tested a well-studied negative odor that activates nociceptors in fish, mustard oil (allyl isothiocyanate), and found that like zebrafish, both surface fish and cavefish exhibited negative odor indices (Fig. 4a&b, yellow label; Supplemental Figure S2, Supplemental Table S15). After establishing a negative odor, we tested three odors derived from food sources, shrimp(39), worms(40) and bat guano(41), that are naturally found in Mexican caves and rivers. Again, both surface fish and cavefish exhibited positive odor indices for all food odors tested (Fig. 4a, green labels; Supplemental Figure S3, Supplementary Table S4, Supplemental Table S16-19). These experiments show that cavefish maintain an ancestral odor perception of aversion to painful odors and approach movements towards food odors.

**Fig. 4.**
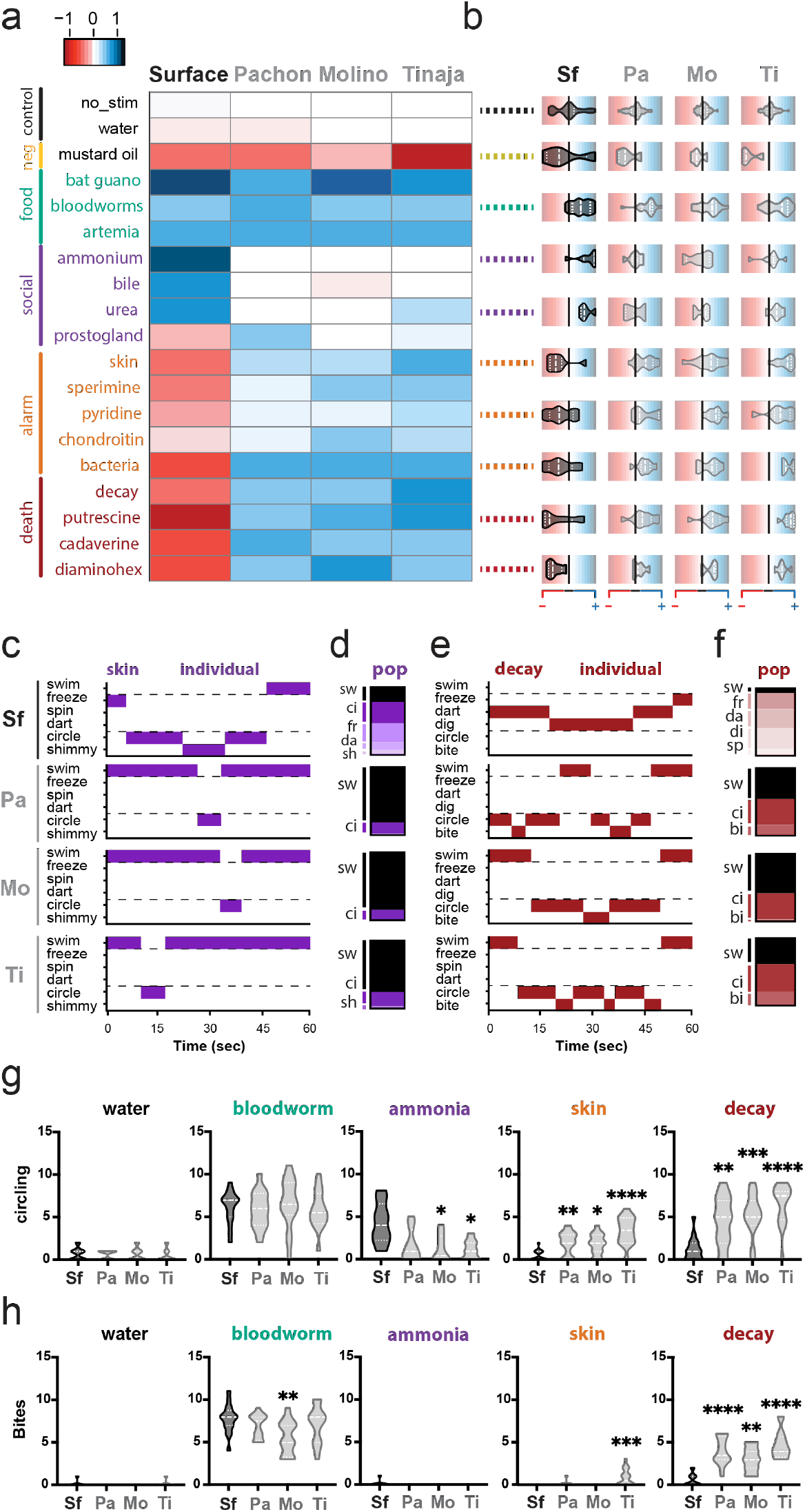
Cavefish exhibit approach behaviors when exposed to ancestrally negative alarm and decay odors. a. Pairwise heatmap of odor index averages between odor and *A. mexicanus* population. Odor indices were encoded as positive (dark blue,1) to negative (dark red, -1) colors to help visualize attraction and repulsion to different odors (sample sizes in Table 3). b. Horizontal violin plots that exhibit positive (blue) and negative (red) odor index values for each population. Each row corresponds to the odor row and color values depicted in the heatmap. c-f. Representative and population ethograms for (cd) social odors and (ef) decay extract behaviors, where each row corresponds to a specific population. gh. Quantification of (g) circling and (h) bite bouts in surface fish and cavefish populations exposed to water, bloodworms, ammonia, skin and decay extract. Odor Indices were compared using non-parametric ANOVA’s, followed by Tukey corrected p-value multiple comparisons. p-values; *-p<0.05*, ***-p<0.01***, -p<0.001.

### Only female cavefish exhibit approach behaviors when exposed to common social odors

Previous work in *A. mexicanus* suggests that blind cavefish lost most social behaviors, including shoaling and schooling, and are considered asocial (24, 42–44). Thus, we wondered whether an asocial organism would respond to common social odorants. To answer whether cavefish maintain ancestral social odor perception, we tested our fish using urea, ammonia, bile, and prostaglandin F_2_*a* that have been shown to elicit kin recognition and breeding in other fish (29-31). We found that surface fish exhibit positive odor indices for ammonia, bile and urea, which suggests these odors are ancestrally positive (Fig. 4a-c, purple label; Supplementary Figure S4, Supplemental Table S20-27). Conversely, we found that cavefish exhibited no variation in behavior when exposed to social odors, suggesting a loss of social odor perception (Fig. 4a-c, purple labels; Supplementary Figure S4, Supplemental Table S20-26). In addition, we found that prostaglandin F_2_*a* produced no change in behavior across all populations (Supplementary Figure S4, Supplemental Table S26). However, when we examined populations by sex, we found that cavefish exhibit sexual dimorphism, with females exhibiting positive odor indices and males showing neutral indices (Supplementary Figure S5, Supplemental Table S2728). These experiments suggest that social odor perception has evolved in cavefish to be sexually dimorphic, with only females exhibiting positive odor indices in response to social odors.

**Table 3.** Sample sizes by odor and fish population.

| **Odor** | **Surface** | **Pachon** | **Molino** | **Tinaja** |
| --- | --- | --- | --- | --- |
| Control (DMSO) | 46 | 45 | 52 | 53 |
| Mustard Oil | 4 | 4 | 4 | 4 |
| Bloodworm | 16 | 16 | 18 | 16 |
| Shrimp | 8 | 8 | 8 | 8 |
| Bat Guano | 8 | 8 | 8 | 8 |
| Taurocholic Acid | 10 | 10 | 10 | 10 |
| Ammonium Chloride | 8 | 8 | 8 | 8 |
| Prostaglandin F2a | 8 | 8 | 8 | 8 |
| Urea | 9 | 9 | 9 | 9 |
| Skin Extract | 19 | 19 | 16 | 16 |
| Pyridine-n-oxide | 10 | 10 | 10 | 10 |
| Chondroitin Sulfate | 8 | 8 | 8 | 8 |
| Bacteria | 8 | 8 | 8 | 8 |
| Decay Extract | 16 | 15 | 15 | 14 |
| Spermine | 10 | 10 | 10 | 10 |
| Cadaverine | 10 | 10 | 10 | 10 |
| Putrescine | 10 | 10 | 10 | 10 |
| Diaminohexane | 10 | 10 | 10 | 10 |

### Cavefish exhibit approach behaviors when exposed to alarm and death associated odors

Finally, alarm and death-associated odors elicit strong alarm responses in zebrafish (11, 12, 28, 29, 31–33, 36). For instance, zebrafish exhibit a range of behaviors when exposed to these odors, including extreme stimuli avoidance towards decay odorants, such as putrescine and cadaverine. Similarly, we found that surface fish on average exhibited negative odor indices when exposed to alarm substances, such as spermine and commensal skin derived bacteria Staphylococcus saprophyticus, and death odors, such as putrescine and cadaverine (Fig. 4a&b, red labels; Supplementary Figure S6, Supplemental Table S29-37). However, ethograms provide a more complex picture, with surface fish showing a range of erratic bottom dwelling, freezing and corner digging (Fig. 4e&f, red labels). Conversely, cavefish exhibited neutral to positive indices, that are characterized primarily by circling behavior (Fig. 4e, red labels; Supplementary Figure S6, Supplemental Table S29-37). Furthermore, all cavefish populations exhibited positive odor indices and increased positive bout types across all death associated odors, including an increase in circling and biting behavior (Fig. 4e&f; Supplementary Figure S7, Supplementary Videos S9, Supplemental Table S38-45). Taken together, these results suggest that cavefish exhibit approach behavior with positive behavioral bouts when exposed to environmentally derived alarm and death-associated odors.

### The range of surface to cavefish *F*_2_ hybrid odor indices suggests that social, alarm and death odor perceptions are genetically inherited

Our odor perception results suggest that cavefish have evolved novel odorant perceptions in response to social, alarm and decay associated odorants. Our next question was whether these neurobehavioral changes are genetically encoded. Although many cave populations are physically separated from surface dwelling populations, surface and cave adults can be bred to produce fertile hybrid offspring (21, 45). Hybridization provides a functional genetic tool to determine the heritability of a trait and facilitate the mapping of genetic variants to biological traits. To test whether variation in odor perception is genetically encoded in cavefish, we crossed surface fish to Tinaja cavefish and backcrossed the offspring to test surface to cave F_2_ hybrids for odor perception behavior (Fig. 5a). We tested each individual F_2_ hybrid for social (ammonia), alarm (skin extract) and death (decay extract) odor perception and then assessed correlations with biological features, such as eyes, sex and behavior-to-behavior relationships (Fig. 5b-i). First, we found that each odorant class elicited a range of odor indices across the hybrid population, suggesting that these behaviors are genetically encoded (Fig. 5b, Supplemental Table S46). Next, while we did not find any associations between anatomical features and behavior (Fig. 5c&d, Supplemental Table S47-53), we did discover that hybrid females exhibited positive odor indices when exposed to social odors, but did not find sexual dimorphism when exposed to skin or decay odors (Fig. 5e&f, Supplemental Table S54-56). Finally, we found that hybrid populations exhibit expected behavior-to-behavior correlations, such as a negative association between social and alarm behaviors, and a strong positive association between alarm and death associated behaviors (Fig. 5g-i, Supplemental Tables S57-59). The observed hybrid odor perception range and associations between behaviors suggests that odor perception differences in surface fish and cavefish are genetically encoded and that similar brain regions may encode variation across similar odor subtypes.

**Fig. 5.**
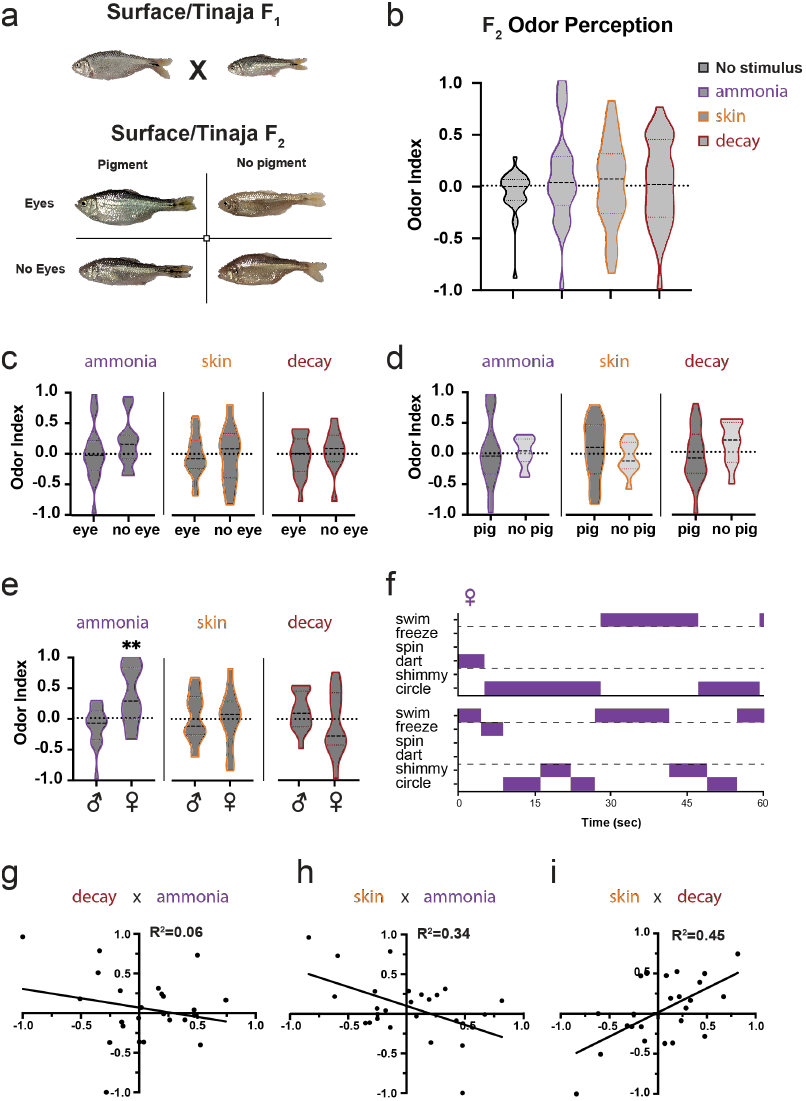
Surface to cave *F*_2_ hybrid behavior suggests olfactory perception is genetically inherited. a. Surface to Tinaja F_2_ hybrids were generated by crossing Surface to Tinaja *F*_1_ siblings. b. Hybrid population behavior for social (purple), alarm (orange) and death (crimson) odors span the full approach to avoidance value range (n=29). c-e. Hybrid population comparisons binned by (c) eyes, (d) pigment and (e) sex (male = 17, female = 12). f. Representative ethograms exhibiting female circling and shimmy bouts that are indicative of mating. g-i Pearson correlations for (g) decay versus ammonia, (h) skin versus ammonia and (i) skin versus decay. Indices were compared using non-parametric ANOVA’s, followed by Tukey corrected p-value multiple comparisons. p-values; *-p<0.05*, ***- p<0.01***, -p<0.001.

### Starvation increases attractive and bold responses from surface fish when exposed to negative odors

Our surface to cave F_2_ hybrid behavioral experiments suggest that odor perception is genetically encoded in surface and cave populations. However, we wondered whether the odor perception behavior we observed is phenotypically plastic and whether food availability impacts avoidance/attraction responses in the ancestral state? To determine the plasticity of odor perception, we withheld food for one-month (30 days) and repeated our olfactory perception assays for skin (skin extract) and death (decay extract) odors at the end of every week (Fig. 6a). First, we found that cavefish exhibited similar indices and bout types across the month of testing, with little variation in valence or group ethograms (Fig. 6b&c, Supplementary Table S60-69). Second, we found that surface fish on average exhibited neutral odor perception indices by week 3 and week 2 for skin extract and decay extract, respectively (Fig. 6b&c). Finally, we found surface fish exhibited positive odor indices and some striking behavior for decay extract at week 3 and 4, and positive odor indices with no feeding related behavior for skin extract in week 4 (Fig. 6d&e). While our sample sizes were modest, these results suggest that odor perception is plastic and that food availability may provide a driving force for odor perception evolution in cavefish.

**Fig. 6.**
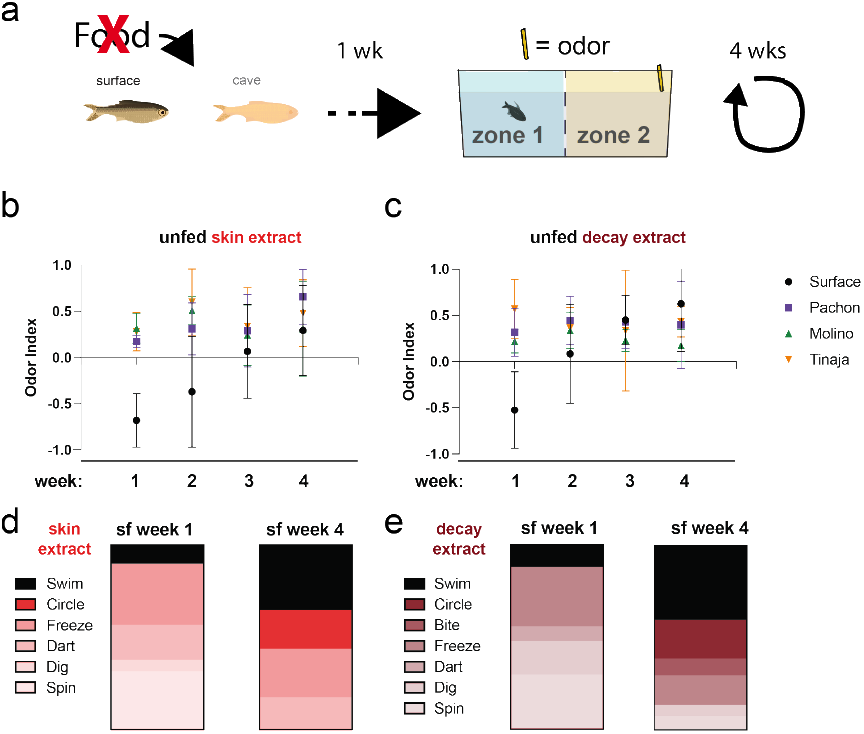
Food availability impacts avoidance/attraction behaviors and hunger drives surface fish towards negative odors. a. Pictograph demonstrating the paradigm for withholding food across 4 weeks and measuring odor perception at the end of each week. bc. Behavioral analysis of odor perception indices for (b) skin extract and (c) decay extract exposed fish across four weeks. de. ethogram charts displaying bout percentages for (d) skin extract and (e) decay extract exposed surface fish at the beginning and end of the experiment. Indices were compared using non-parametric ANOVA’s, followed by Tukey corrected p-value multiple comparisons. p-values; *- p<0.05*, ***- p<0.01***, - p<0.001.

### Brain activity mapping finds overlap in activity of the thalamus and preoptic region of food and decay odor exposed cavefish

While cavefish exhibit positive odor indices and swimming bouts when exposed to a range of odor classes, we wondered whether these different classes of odors elicit similarities in neural activity across the forebrain. The protein total-ERK (t-ERK) becomes phosphorylated (p-ERK) following an action potential and p-ERK expression in active neurons peaks within 3 minutes of firing (46). Therefore, this phosphorylation event provides a minute-scale biomarker for neural activity that is perfect for mapping responses to sensory stimuli. To map odorant stimulated neural activity, we first exposed individual Pachon cavefish to one odorant for 10 minutes before whole-brain protein staining and tissue clearing (Fig. 7a). Then, to analyze across samples, we generated an autofluorescence template and reference brain to register all brain images into the same 3D space (Fig. 7a-c). Finally, we used conspicuous features of the tERK expression pattern (e.g. structural) and food exposed p-ERK expression patterns (e.g. physiological) to segment identifiable subregions that are likely to correlate with odor attraction and appetite (Fig. 7c&d). We therefore focused our analysis on subregions of the dorsal pallium (amygdala-like), thalamus and preoptic region (Fig. 7d). We found that food exposed, and decay exposed fish exhibited similar increases in p-ERK levels of the thalamus and preoptic region, while skin and decay exposed fish showed similar increases of p-ERK levels of the dorsal pallium (Fig. 7e&f, Supplemental Tables S70-76). Of these four regions, the thalamus exhibited the largest numerical and most visibly discernable variation across conditions, with bilateral neuropil regions exhibiting the same increase in pERK across food and death odor exposed cavefish (Figure 7e; flanked by yellow asterisks). While our sample sizes are low, these activity patterns suggest that decay and food scents elicit similar signatures within regions known to contribute to appetite and odor perception.

**Fig. 7.**
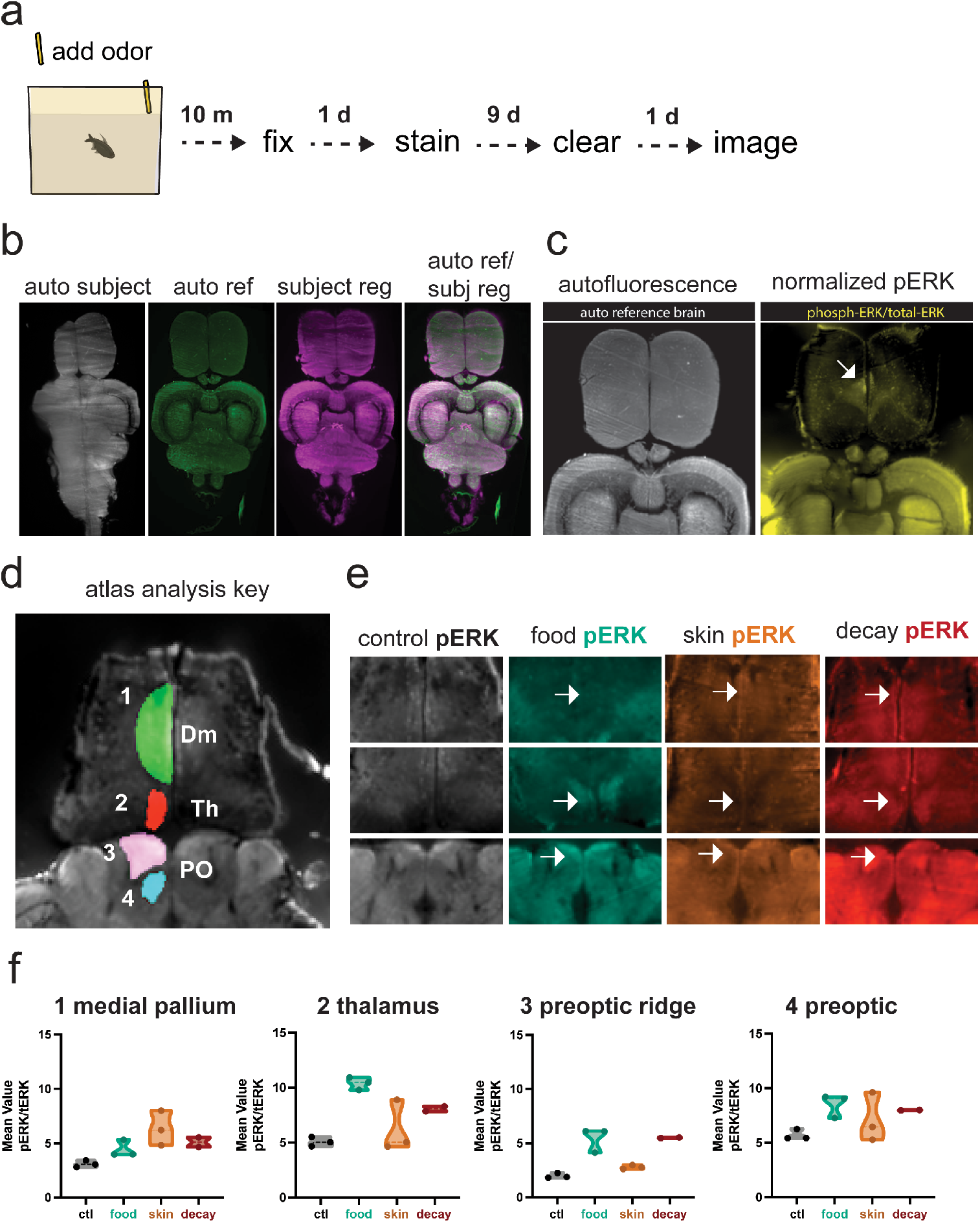
Brain mapping in the forebrain finds similar activity patterns in the cavefish thalamus and preoptic region when smelling food or decay extract. a. schematic of experimental pipeline through two weeks of dissection, staining, clearing and imaging b. example of adult brain registration where autofluorescence (auto subject) is used to register to our reference brain (auto ref). The final product is a reformatted brain (subject reg) in the reference space (auto ref/ subj reg). c. example of normalized pERK in the pallium of a skin exposed fish (arrow pointing to dorsal pallium, e.g. amygdala correlate). d. atlas analysis key showing the four subregions that were analyzed for activity mapping. Dm, medial pallium; PO, preoptic region; Th, thalamus. e. Optical sections displaying normalized p-ERK signal in Pachon cavefish exposed to DMSO (control p-ERK), bloodworm water (food p-ERK), skin extract (p-ERK) and decay extract (decay p-ERK). Each column of images is derived from a single fish for each condition, top anterior to bottom posterior positions. Flat arrows point to pallial and preoptic regions of activity, while yellow asterisks flank high p-ERK in the thalamus of food and decay exposed fish f. Box plots comparing regional p-ERK intensities of Pachon cavefish exposed to four conditions, control (DMSO), food (bloodworm water), skin (skin extract) and decay (decay extract). Indices were compared using non-parametric ANOVA’s, followed by Tukey corrected p-value multiple comparisons. p-values; *- p<0.05*, ***- p<0.01***, - p<0.001.

## Discussion

Odor perception and odor induced behaviors allow organisms to survive by pursuing food and mates, while avoiding predation and disease. The odor perception of most organisms is conserved across ecological niches, with positive valences and approach toward food and social odors, and negative valences and avoidance towards threats of predation and death (7, 8, 47, 48). Our odor perception screen suggests that cavefish olfactory behavior has changed from avoidant and negative to approach and positive in relation to signals of predation and death. The frequency and magnitude of these changes likely reflect the change in evolutionary pressures experienced in cave pool environments, such as a reduction in nutritional availability and decreased predation. This initial behavioral study provides a new neurogenetic model for uncovering the genes and neurons that functionally control odor perception in vertebrates.

### Approaching dead and dying fish may improve resource allocation in a food variable environment

This study motivates a straightforward question: why do cavefish exhibit approach behaviors when exposed to odor sources that could be dangerous or deadly? We suggest that strong selective pressures via reduced predation or lack of resources caused a change in odor perception from odor avoidance to odor approach. Indeed, we know that cavefish have been found to be opportunistic omnivores, with gut contents ranging from crustaceans to other cavefish, and some mostly consisting of bat guano (39, 49). For instance, cave fry from the Pachon cavefish feed mostly on crustaceans, while adult guts contain mostly detritus (39, 50). However, we do not have enough information about the seasonality and abundance of these food sources across different cave localities. Therefore, a rewiring of odor perception for ancestrally negative odors would provide a new foraging strategy for acquiring any possible food source. Indeed, most carrion obligate avians use decay odors to find carcasses, but omnivorous predators may also scavenge using the same odors when resources are dire (51, 52). Furthermore, our finding that cavefish exhibit more positive behavior towards odors of decay versus alarm associated odors, may suggest that an injured but reactive fish is less attractive than a lifeless decaying fish, which would not present an immediate threat (albeit a possible immunological risk by spreading disease (53)).

### De novo mutations or genetic assimilation may underlie the switch from avoidance to approach for alarm and death odors

Our results also present a fundamental question of evolution: how does a new phenotype come to exist? Work across many taxa suggests that phenotypes can arise due to either de novo genetic mutations or plasticity, when environmental factors favor phenotypes from existing or ancestral alleles (54). For instance, De novo mutations in several cavefish populations have been shown to influence novel phenotypes, such as eye and pigment loss (55, 56). In regards to our study, selection may act upon de novo genetic variants in cavefish to rewire odor perception due to selective pressures in the cave environment, such as availability of food (54). Alternatively, cavefish researchers have shown that phenotypic buffering can mask cave-like traits in surface fish due to canalization. For example, researchers revealed the presence of a small-eye size phenotype in surface fish by treating developing fish with an hsp-90 inhibitor (57). Hsp-90 inhibition of developing surface fish caused a wide variation in eye size that was shown to be transgenerationally inherited by backcrossing surface fish with small eyes. More recently this phenomenon was investigated in spadefoot tadpoles to determine how genes and environmental factors impact the emergence of cannibalism (58). In that study, researchers compared cannibalistic versus non-cannibalistic genera and found that changes in the environment, such as space and resource allocation, produced the emergence of cannibalism in the non-cannibalistic genera. Additionally, they found that genetic variation played a role in the emergence of cannibalistic behavior that suggests ancestral alleles provide phenotypic plasticity for reducing competition in a crowded environment (58). Therefore, surface fish alleles related to odor perception may provide evolutionary capacitance for attraction to negative odors that is suppressed or masked in ideal conditions. This phenomenon would explain why unfed surface fish exhibit an increase in odor indices and positive swim bouts when exposed to death odors.

### Hunger driven changes in surface fish behavior suggests that odor perception plasticity may play a role in the evolution of foraging

Previous research has shown that the availability of food impacts avoidance and attraction in response to odorants and tastants (59–61). We found that 3-4 weeks of starvation resulted in positive bout types, including striking behavior in surface fish when exposed to decay odors. This suggests that hunger impacts the reception or integration of negatively associated odors in fish. Indeed, research in Drosophila have shown that starved flies are bolder in their approach to negative odors/tastants that are deadly to ingest (59). For example, the sensitivity and perception of odors change in response to starvation, with increased neuropeptide F neuron sensitization to attractive odors and concomitant suppression of aversion via tachykinin signaling (60, 62). Odor perception plasticity has also been examined in C. elegans, with starvation impacting odor discrimination and enhanced odor memory in relation to novel odors (10, 14). Therefore, these studies provide examples of neural plasticity that could explain starvation induced olfactory perception behavior observed in surface fish. We will further investigate the heritability of ‘bold olfactory perception’ in surface fish by backcrossing these bold surface fish and testing odor perception and starvation induced odor perception in F_1_ bold surface offspring (see previous section).

### Sexual dimorphism in relation to social cues suggests a need for females to find males

Social cues can provide fitness advantages for individuals within a population through social coherence and mating (63). Surprisingly we found that only female cavefish exhibit positive odor indices when exposed to social odors that are positive for both male and female surface fish. Previous fish studies have described sexual dimorphism in relation to olfactory ability, including other Characids (64), and variation in olfactory organs anatomy and physiology, such as in deep-sea fish (65, 66). Larger nostrils, olfactory lamellae and enhanced physiology was associated with males of these species, suggesting that males needed larger and more sensitive olfactory organs for finding female mates in sparsely populated areas of the deep ocean (54). Therefore, cavefish females may have retained approach to social cues to find males for mating in cave habitats. Indeed, we found convergence across cave populations and a strong social odor approach phenotype in F_2_ hybrid females, suggesting that this trait could be sex-linked via sex chromosomes.

### Hybrid range of odor indices suggests that cavefish odor perception is a convergent cave adapted trait that arises from standing genetic variation

While our feeding experiments support the role of ancestral alleles, our hybrid experiments suggest that genetic variation across populations drives odor perception. All three cave adapted populations of *A. mexicanus* tested here exhibited the same changes in odor perception, including similar bout types and bout frequency, when exposed to our four death-associated odors. The range of behaviors and bout types for both surface and cave fish was then recapitulated across the F_2_ hybrid population. These experiments suggest that changes in odor perception are genetically encoded, which could include genes impacting the nervous system (67, 68), immune system or metabolism (3, 69, 70). For instance, research on scavengers suggests that behavioral evolution towards searching for carrion includes expansions of olfactory receptors, or expression/coding of digestive enzymes and immunological pathway proteins (51, 52, 69). While we have yet to perform a QTL mapping experiment, previous cavefish trait association studies have found that scavenger linked genes, such as the immune cell regulator bcl6a and the cell adhesion molecule ep-cam, are under selection in different lineages of cavefish (71). Our future studies will utilize QTL mapping to investigate the evolutionary genetics of olfactory perception in vertebrates

### Food and decay derived odors elicit similar brain activity signatures that suggest may underlie similarity in behavioral repertoires

Our adult forebrain activity mapping found that brains exposed to food and decay extract had similar activity profiles in the thalamus and preoptic region. This is especially true for large bilateral neuropil regions of the thalamus (Figure 6c; starred panel), where we did not observe large increases in activity when cavefish were exposed to DMSO or skin extract. Previous work in humans and rodents have identified the mediodorsal thalamic nucleus (MDT) as a sensory integrator that plays important roles in odor discrimination and perception (72, 73). For instance, ischemic damage to the medial-dorsal thalamus can influence perception, resulting in parosmia like deficits in which the pleasant smell of food becomes nauseating (74, 75). Similarly, researchers have shown that the MDT plays an important role in discriminating different odor mixtures, with deactivation of the region causing animals to reduce consumption of preferred odor-taste mixtures (76). In addition, we also see similar activity patterns in subregions of the preoptic region, which may represent a novel role for odor perception outside of the preoptic regions well known role for processing social odor cues (77, 78). While our dataset is small and spatially biased, the overlap of activity between food and decay exposed brains suggests that future work will allow us to map olfactory perception to specific regions and cellular subtypes.

## Conclusions

In conclusion, this study describes a convergent change in cavefish odor perception that causes cavefish to respond positively to ancestrally negative odor cues. These behaviors were shown to be convergent across all cave populations tested, hybrid experiments suggest these behaviors are genetically encoded, starvation influences behavioral plasticity and neurological correlates suggest multiple brain regions drive perception differences based on odorant class. Overall, our work provides a novel model for investigating the evolution of complex phenotypes and the mechanisms driving odor perception in vertebrates.

## Supporting information

statistical_tables

## ABOUT THIS MANUSCRIPT

This manuscript was prepared using R_*χ*_iv-Maker v1.18.3 (79).

## DATA AVAILABILITY

All statistical tables and custom codes can be found at olfaction. All raw and analyzed data will be made public upon publication.

## ACKNOWLEDGEMENTS

We would like to thank the administration and staff in the Department of Biology at St. John’s University and in the Department of Anatomy and Neurobiology at the University of Puerto Rico School of Medicine.

## Methods

### Animal care and husbandry

All populations are kept in housing tanks in the *A. mexicanus* fish facility located in the Department of Biological Sciences at St. Johns University. *Astyanax mexicanus* stocks are maintained and cared for according to the St. John’s University IACUC approved protocol 2073. Adult fish are maintained on a recirculating z-hub system (Pentair Inc., Minneapolis, MN, USA) and custom-built recirculating racks with the following parameters; temperature 22-23 °C, conductivity 200-900 *µ*S and pH 6.8-7.8. Fish are maintained on a 14:10 ZT light:dark schedule. Fish stocks include wildtype: Surface fish Rio Choy, Pachon cavefish, Molino cavefish, Tinaja cavefish and hybrids: Surface/Tinaja *F*_1_ and F_2_ hybrids.

### Odorant preparations

Commercially purchased odors were measured and dissolved in system water with 0.5% DMSO (Table 1). DMSO was added to each solution as a molecular weight for delaying odorant dispersal across odor zones (Supplementary Figure S5). All solutions were dispensed as 50 uL aliquots into 9-L behavioral tanks. Bloodworm water was created by soaking 1 gram of frozen bloodworms in 25 mL of system water at 4 °C overnight. This solution was then centrifuge and the aqueous phased extracted as the odorant. Shrimp water was derived by soaking 1 mL of concentrated artemia shrimp in 25 mL of system water at 4 °C overnight. This solution was then centrifuge and the aqueous phased extracted as the odorant. Bat guano water was made by soaking 1 gram of bat guano fertilizer in 1 mL of system water and heated at 95 °C for 1 hour. This solution was then centrifuge and the aqueous phased extracted as the odorant. Skin extract was prepared by euthanizing a surface fish or cavefish in 500mg/L Tricaine-S (MS222), scoring the skin with a scalpel and then rocking the scored fish in 40 mL of system water for two hours. Immediately following, skin extract liquid was then aliquoted into 1.5 mL tubes and heated to 95 °C for 30 minutes before being stored at -80 °C for up to 1 month. Skin extract was then diluted 1:10 in 0.5% DMSO and system water as a working stock. Decay extract was derived using skin extract fish by adding 40 mL of system water and rocking overnight at room temperature. This decay extract water was treated the same as the skin extract (heated and spun) and stored at -80 °C for up to 1 month. Bacteria odor was derived by preparing commensal strain Staphylococcus saprophyticus ATCC 15305 extracts grown in BH infusion medium (Brain Heart Infusion Broth media BD237500; BD, Franklin Lakes, NJ). This protocol was adapted from a previous odor study in zebrafish (32). 0.5 mL of an overnight pre-culture were used to inoculate 50 mL of fresh media. After a 3-hour incubation (30 °C, 220 rpm; New Brunswick orbital shaker; Eppendorf, Hamburg, Germany) cells were harvested by centrifugation (3000g; Eppendorf 5804R centrifuge, Hamburg, Germany) and the pellets stored at -80 °C for up to 1 month. To prepare the extract, one pellet corresponding to 25 ml of original culture was resuspended in 2.5 mL of 0.7% NaCl, and then washed twice with 2.5 ml of H2O. The resulting resuspension was incubated for 30 minutes at 90 degreees in a water bath to lyse bacteria. Cell debris was removed by centrifugation while the supernatant was recovered as the odorant preparation.

### Video capture and analysis

Behavior was captured using an acA1300-200um c-mount camera (Basler, inc., Exton, PA, USA), with a 16mm C Series Lens VIS-NIR (Edmund Optics, Barrington, NJ, USA) via the Pylon software (Basler, Inc.) at a frame rate of 25 fps. Each video was then initially analyzed using the Ethovision XT-14 software (Noldus, Inc, Leesburg, VA, USA). Ethovision tracking was performed using the dynamic subtraction setting set to subject lighter and darker. Tracks were then time binned and zone time cumulative distance was exported to derive the odor indices.

### Olfactory behavioral experiments

Fish were transferred from the fish facility to the animal behavior room in groups of eight. They were then transferred individually to 9-liter behavioral tanks (Iwaki, Inc.; Holliston, MA, USA) and acclimated to the behavior room overnight. Tanks were then illuminated by either direct diffused light (200-220 lux), indirect white light (<0.1 lux) or direct diffused infrared light (180-240 IR lux). Experimenters used custom made cellphone night vision goggles for direct diffused infrared light experiments in the dark. DIY night-vision goggles consisted of an infrared camera (FLIR ONE Generation 3 Thermal imaging camera for Android, Teledyne, Wilsonville, OR, USA) and modified Google Cardboard (Fannego, Amazon, Inc., Seattle, WA, USA) affixed to laboratory goggles. Each odorant trial included two 5-minute videos, a pre-stimulus control video (no stimulus) and an odor exposed video (e.g. bloodworm). To test olfactory perception, 50 *µ*L of odorant was added to a randomized size of the tank and behavior was captured for 5 minutes. In-between odor trials, fish were removed from behavior tanks and kept in 2.5-liter zebrafish tanks while experimenters cleaned each behavior tank with 70% ethanol and hot water. Fish were then placed back in behavior tanks and given 2-hour to acclimate before subsequent odor trials. Fish were subject to no more than 2 odors per day, to reduce stress and olfactory overstimulation. Following experiments, adults were transferred back to the fish facility and returned to housing tanks.

### Approach and avoidance analysis

Behavioral videos were analyzed using the Ethovision XT-16 software (Noldus, Inc., MA, USA). Ethovision tracking was performed using the dynamic subtraction setting set to subject darker, and cumulative time in zone 1 (no odor) and zone 2 (odor) were used to calculate odor indices.

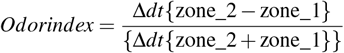

Initially, behavioral videos for surface fish were binned at 1-minute intervals to determine peak approach for bloodworm water (significant increase in odor indices). The peak of 1-minute was then used to bin and analyze all further odorants tested in surface fish and cavefish. Numbers that are positive indicate approach (more time in zone 2; odor zone), while negative numbers indicate avoidance (more time in zone 1; no odor zone). In addition, swimming velocities were automatically calculated using the EthoVision XT software, while fish bouts and poses were called and tallied by trained investigators Ethograms were then constructed using custom python scripts and exported as pdfs.

### Surface to Tinaja cavefish *F*_2_ hybrid olfactory behavior and correlations

Individual Surface to Tinaja F_2_ hybrid fish were assayed for ammonium chloride, skin extract and decay odor. Fish were behaviorally assayed as previously described and then kept in a post-assayed tank while the remaining individuals were assayed. Following our behavioral experiments, fish were anesthetized using 500mg/L of buffered Tricaine-S in system water, photographed for anatomical features, including presence and absence of eyes, eye size, presence and absence of pigment, standard length and sex (anal fin spine test and later confirmed with dissection), before fin clipping to preserve DNA samples for future studies. Finally, eye, pigment and sex comparisons were made in a group-wise fashion using odor indices, while a simple linear regression and R-squared values were used to analyze the behavioral relationship between classes of odorants. All data analysis and statistical tests were run using PRISM (GraphPad, Inc.)

### Adult brain dissection, staining and tissue clearing

Pachon cavefish adults (2.6-2.9 cm SL) were added individually to 2.5 liter zebrafish with false bottoms. Fish were then transferred to the behavioral room and left to acclimate for an hour in indirect white lighting. 50 *µ*L of odorant was then pipetted into the tank and fish were exposed to that odor for 10 minutes. Fish were then euthanized by rapid cooling in ice water for 15 minutes, followed by immediate decapitation. Heads were then fixed whole in 1 M NaOH buffered 4% PFA overnight, washed and then dehydrated in 100% MeOH. After 24 hours at -20 °C, fish brains were dissected from the head and rehydrated with 20% steps of MeOH/PBS before being permeabilized in 1xPBS/0.3%TritonX-100/20%DMSO for 2-days at 37 °C with rocking. Brains were then incubated in blocking solution (1xPBS/0.3% TritonX-100/10% BSA/10% DMSO) for 2-days, primary antibody solution (1xPBS/0.3% TritonX-100/5% BSA/10% DMSO/ 1:250 antibody) for 3-days and secondary antibody solution (1xPBS/0.3% TritonX-100/5% BSA/10% DMSO/ 1:500 antibody) for 3-days. Stained brains were then cleared using the CytoVista Tissue Clearing Kit (Thermo Scientific, Waltham, MA, USA).

### Adult brain imaging

Stained brains were mounted to a metal stage using UV light curing resin and imaged using the Miltenyi UltraMicroscope Blaze light sheet microscope (Miltenyi Biotec, Bergisch Gladbach, North Rhine-Westphalia, GER). Image stacks were then captured using the 5x dipping objective at 2x zoom with a resolution of 0.9x0.9x2 microns. Following image capture, brain stacks were down sampled to 7.2x7.2x2 microns and hemibrains were stitched using a custom Fiji script.

### Adult brain template generation

All template construction, brain registrations and segmentation was done using Advanced Normalized Tools (ANTs) which utilizes the snap imaging tool kit (ITK) library (80). Standard brain templates for autofluorescence was generated by applying the antsTemplateConstruction2.sh script to all whole adult brain imaging stacks using minimal user input (see custom ANTs scripts) (81). We then used our initial template target brain as the t-ERK reference image brain to create our analysis atlas by segmenting regions that were visually identifiable by either structural (t-ERK) or physiological (p-ERK) signals in food exposed brains. In addition, updated versions of the adult zebrafish atlas (Wullimann and Mueller, 2016(82); Kenney et al. 2021(83)) were used to identify each region and subregion. Segmentation was performed using the segmentation editor plugin in Fiji (ImageJ, (84)). Each brain was then registered via autofluorescence channels and inverse registered to segment the forebrain for p-ERK expression analysis. For p-ERK expression analysis, each unregistered p-ERK was normalized by dividing p-ERK signal by t-ERK signal using the image calculator function in Fiji. Normalized images were then merged with brain segmentations, before measuring average intensity for each regional segment. Normalized intensity each segment volume was then calculated by importing the normalized p-ERK and inverse registered segmentation stacks for each individual brain into Napari as separate layers (85). Average intensity was then extracted by using the n-simpleITK (measurements) tool.

### Statistical analyses

All statistical analyses were carried out using the statistical software PRISM (Graphpad, inc., USA). Individual odors were analyzed using non-parametric one-way ANOVA’s, and when statistically significant, analyzed using a Tukey p-value adjusted multiple comparison test. Surface fish were treated as the control condition (i.e. ancestral behavior) and compared to each individual cave population via the Tukey-corrected multiple comparison test.

## Supplementary Information

**Sup. Figure S1.**
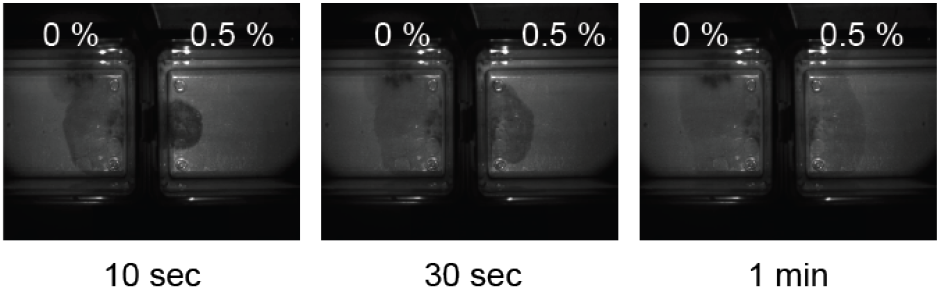
Time series depicting red dye in behavioral tank with and without DMSO. Each image shows red dye with no DMSO (0%) versus red dye with DMSO (0.5%) across three time points, 10 seconds, 30 seconds and 1 minute.

**Sup. Figure S2.**
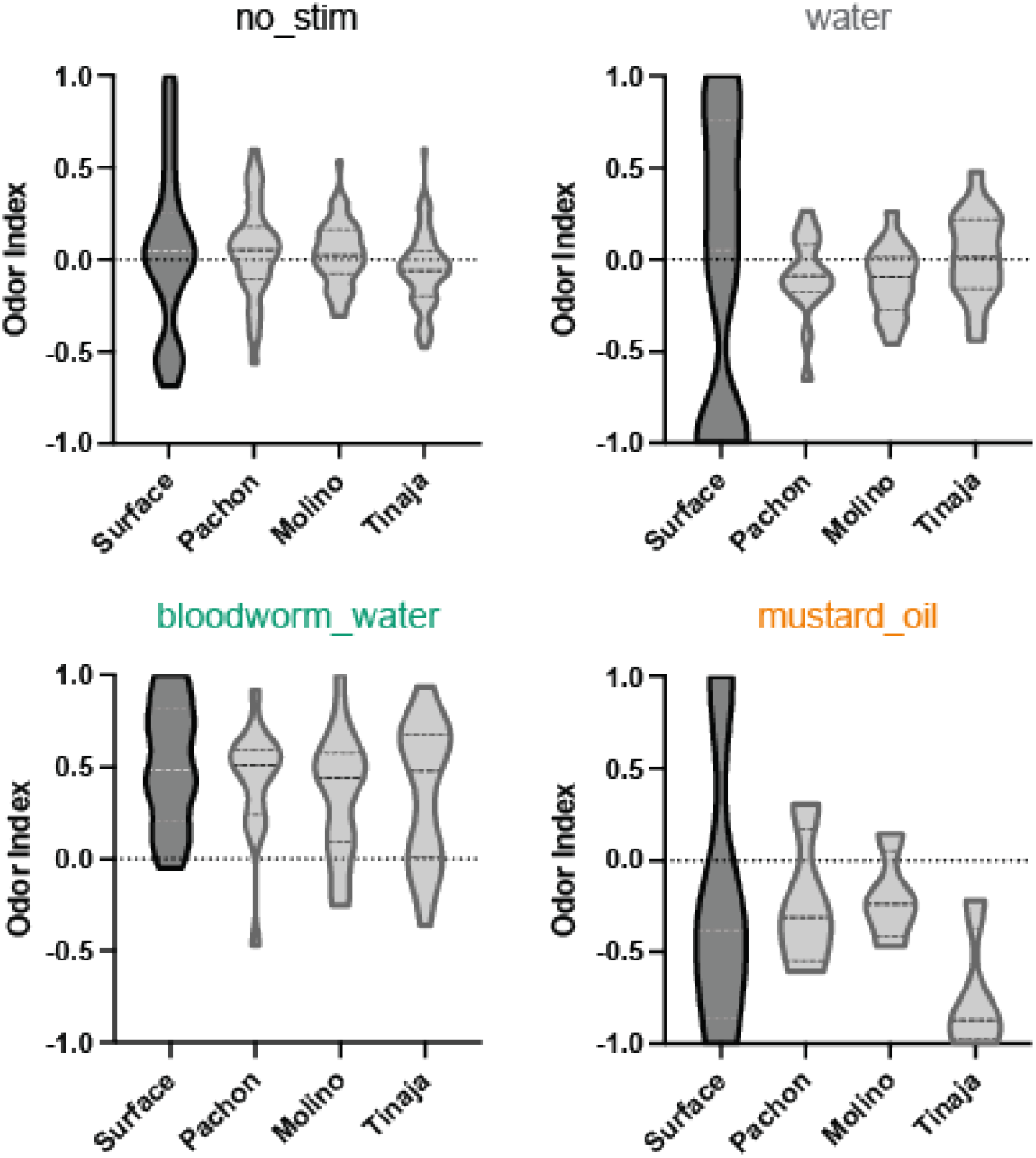
Odor indices for the four control odors. No stimulus = baseline movement, water = odor sham (pipette current), bloodworm water = positive (food) and mustard oil = negative (pain). Each graph represents 1 minute of behavior following the introduction of the title odor.

**Sup. Figure S3.**
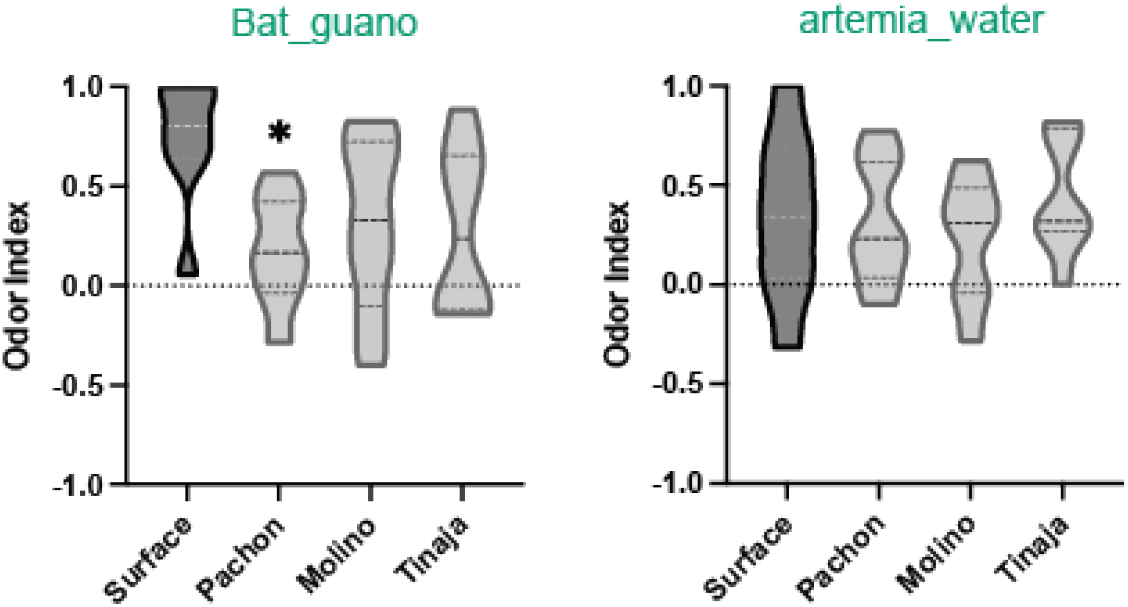
Food odor indices for all populations. Bloodworms are in the previous graph (S3) as an established positive control group. Each graph represents 1 minute of behavior following the introduction of either bat guano or artemia water.

**Sup. Figure S4.**
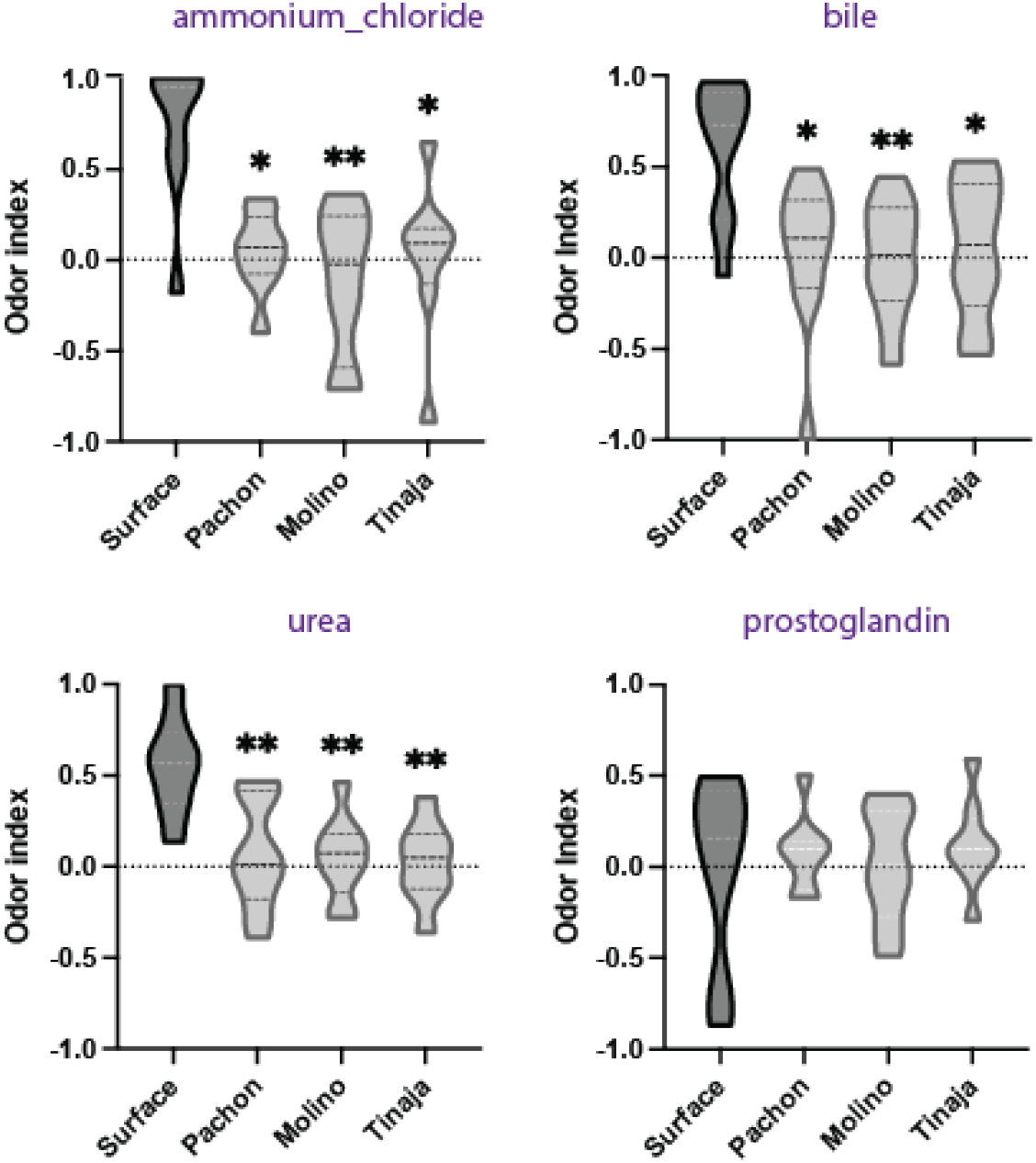
Social odor indices for all populations. Each graph represents 1 minute of behavior following the introduction of either ammonium chloride, bile, urea or prostaglandin F2-alpha.

**Sup. Figure S5.**
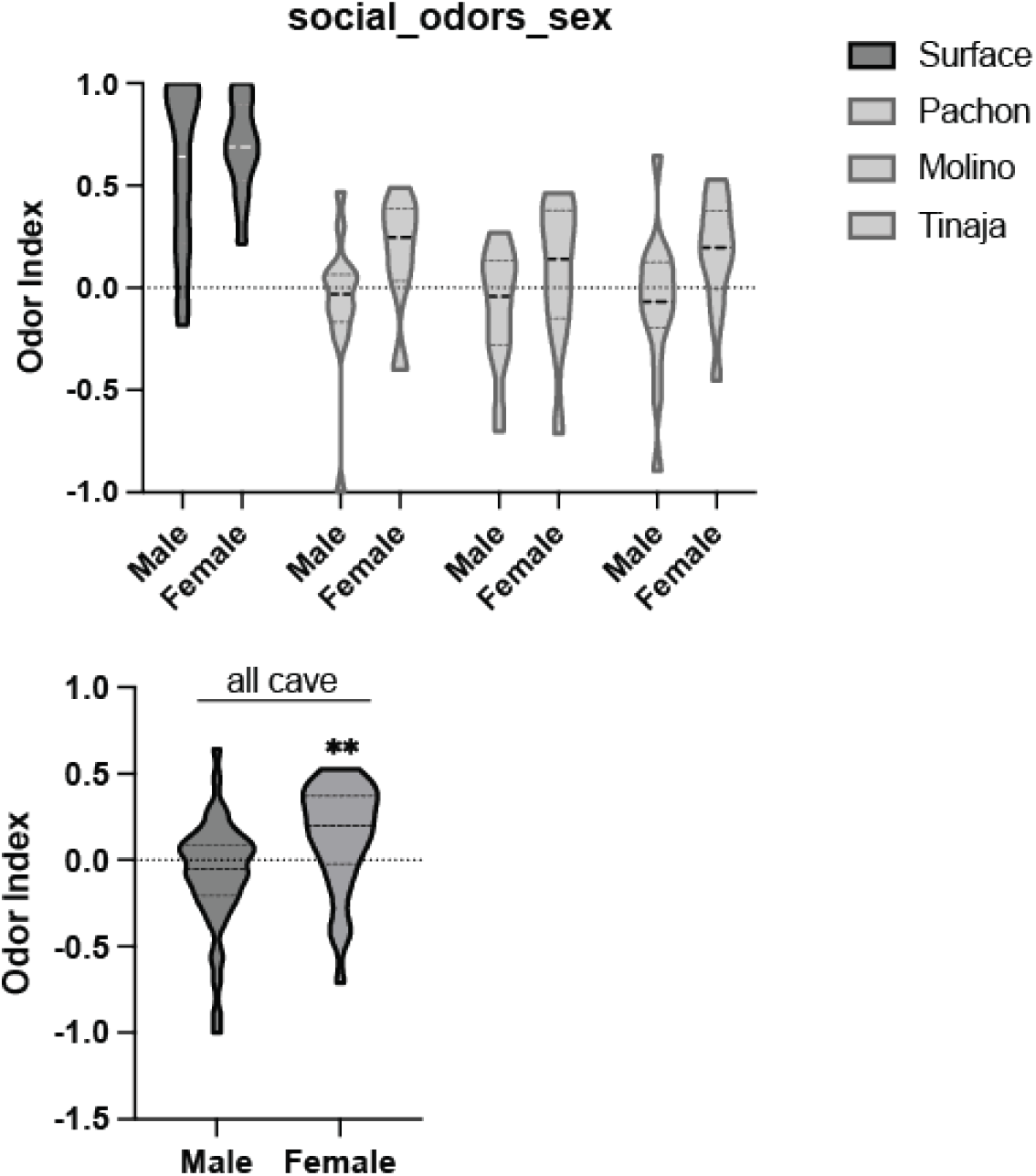
Aggregated odor indices of all social odors for each population. All social data was reanalyzed by sex type and paired statistical comparisons.

**Sup. Figure S6.**
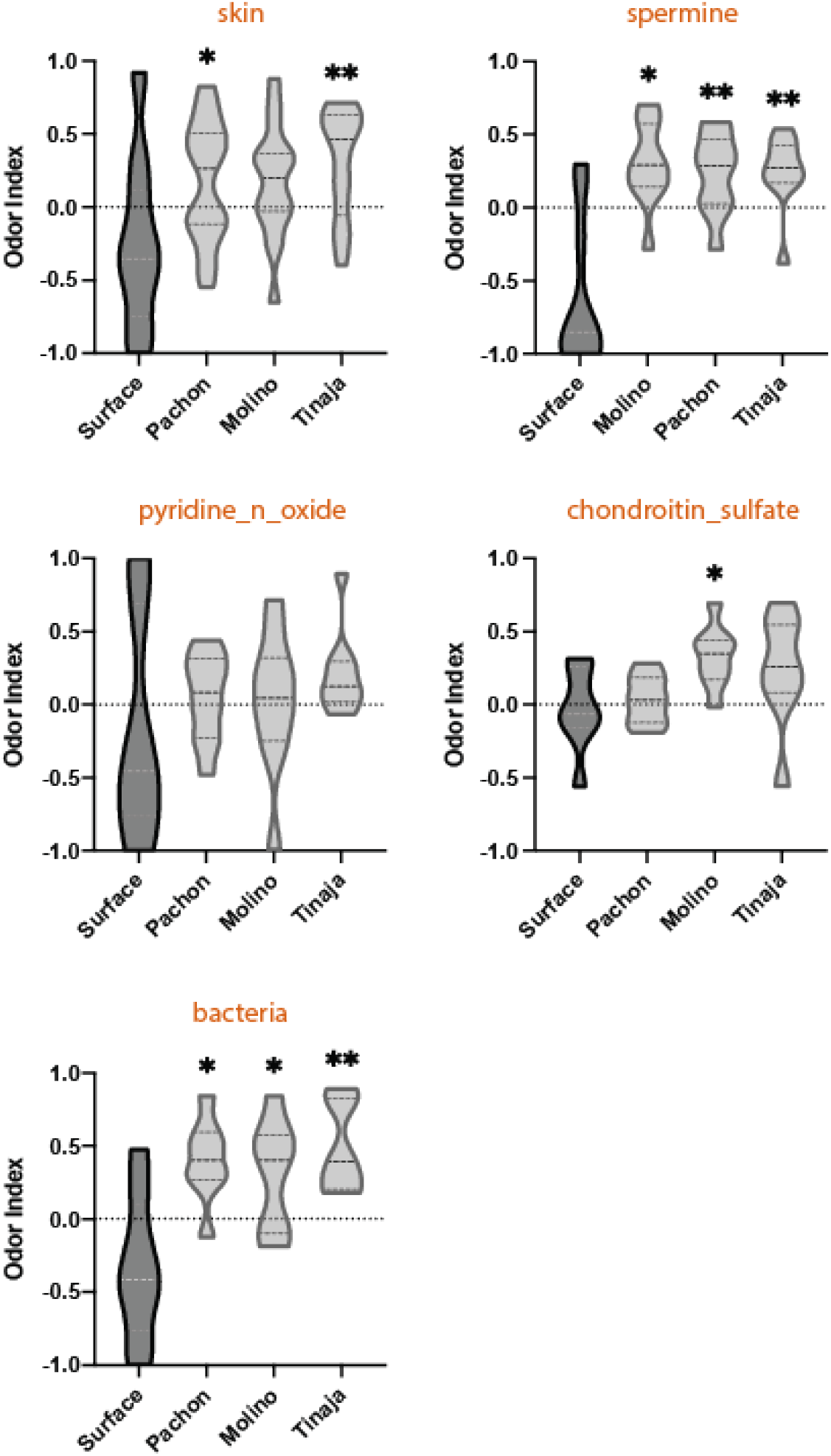
Alarm odor indices for all populations. Each graph represents 1 minute of behavior following the introduction of either skin extract, spermine, pyridine-n-oxide, chondroitin sulfate or bacteria (Staphylococcus saprophyticus).

**Sup. Figure S7.**
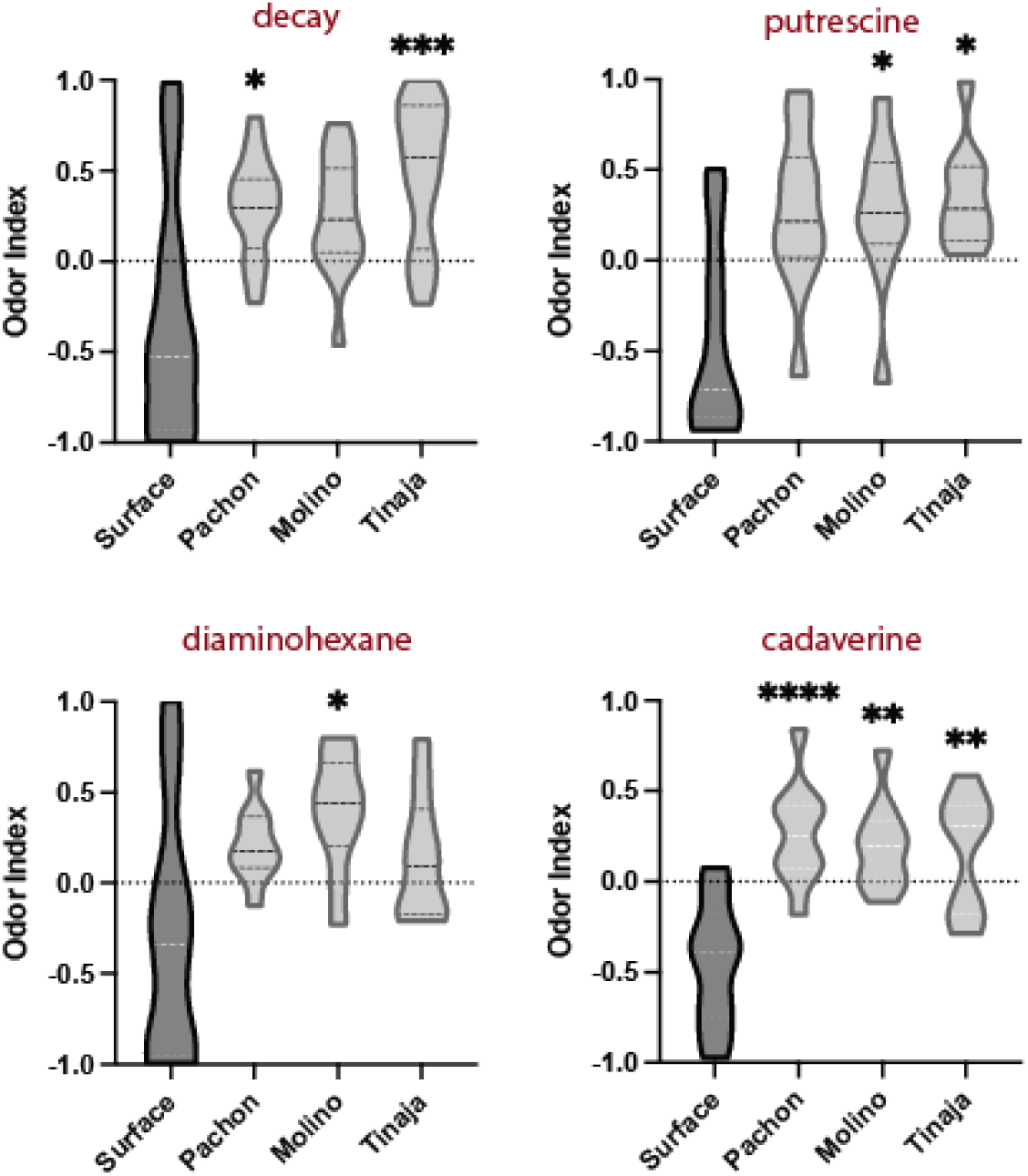
Death odor indices for all populations. Each graph represents 1 minute of behavior following the introduction of either decay extract, putrescine, diaminohexane, or cadaverine.

